# Novel bisubstrate inhibitors for protein N-terminal acetyltransferase D

**DOI:** 10.1101/2021.01.24.427995

**Authors:** Youchao Deng, Sunbin Deng, Yi-Hsun Ho, Sarah M. Gardner, Zhi Huang, Ronen Marmorstein, Rong Huang

**Affiliations:** Department of Medicinal Chemistry and Molecular Pharmacology, Purdue Institute for Drug Discovery, Purdue University Center for Cancer Research, Purdue University, West Lafayette, Indiana 47907, United States; Department of Chemistry, University of Pennsylvania, Philadelphia, Pennsylvania 19104, United States; Department of Biochemistry and Biophysics, Abramson Family Cancer Research Institute, Graduate Group in Biochemistry and Molecular Biophysics, Perelman School of Medicine, University of Pennsylvania, Philadelphia, Pennsylvania 19104, United States

## Abstract

Protein N-terminal acetyltransferase D (NatD, NAA40) that specifically acetylates the alpha-N-terminus of histone H4 and H2A has been implicated in various diseases, but no inhibitor has been reported for this important enzyme. Based on the acetyl transfer mechanism of NatD, we designed and prepared a series of highly potent NatD bisubstrate inhibitors by covalently linking coenzyme A to different peptide substrates via an acetyl or propionyl spacer. The most potent bisubstrate inhibitor displayed an apparent *K*i value of 1.0 nM. Biochemical studies indicated that bisubstrate inhibitor is competitive to the peptide substrate and noncompetitive to the cofactor, suggesting NatD undergoes an ordered Bi-Bi mechanism. We also demonstrated that these inhibitors are highly specific towards NatD, displaying about 1,000-fold selectivity over other closely related acetyltransferases. High-resolution crystal structures of NatD bound to two of these inhibitors revealed the molecular basis for their selectivity and inhibition mechanism, providing a rational path for future inhibitor development.

## INTRODUCTION

α-N-terminal acetylation (Nα-acetylation) is a ubiquitous protein modification that occurs on 80–90% of human proteins ^1^. It is essential for various biological functions, including protein-protein interactions, protein complex formation, cellular apoptosis, rDNA transcriptional regulation, protein subcellular localization, and degradation ^2–4^. This modification is catalyzed by protein N-terminal acetyltransferases (NATs) that transfer an acetyl group from the donor acetyl coenzyme A (AcCoA) onto the Nα-amino group of protein substrates. To date, eight members of eukaryotic NATs (NatA–NatH) have been reported ^5^. Among them, NatA and NatD acetylate the nascent chain after the initiator methionine is cleaved ^6^. NatA is a multisubunit enzyme, acetylating ~40% of human proteins that contain small and uncharged first residues ^1,6^. In contrast, NatD (NAA40) is a monomeric protein showing extremely high substrate specificity for histone proteins H2A and H4 that have the same N-terminal sequence SGRGK ^7,8^.

Nα-acetylation on H4 has diverse biological functions and implications in tumorigenesis ^2,9,7^. Depletion of NatD induces apoptosis through the mitochondrial pathway in colorectal cancer cells ^10^. In addition, NatD is downregulated in hepatocellular carcinoma tissues and upregulated in primary human lung cancer tissues ^11^. As the function of histone H4 Nα-acetylation has recently come to light, it has been shown to regulate crosstalk with arginine methylation, lysine acetylation, and serine phosphorylation on H4 ^12,13^. For example, Nα-acetylation of H4 stimulates ribosomal DNA expression by inhibiting asymmetric dimethylation of Arg3 on H4 (H4R3me2a), consistent with its critical role in cell growth ^12,13^. Nα-acetylation of H4 suppresses Ser1 phosphorylation and induces the expression of Slug transcription to promote the epithelial-to-mesenchymal transition in lung cancer, suggesting that the acetyltransferase activity of NatD is critical for Slug regulation ^13^. In addition, Nα-acetylation of H4 promotes the expression of oncogenes through upregulation of protein arginine methyltransferase 5 in colorectal cancer cells ^10^. Based on these disease connections, NatD has surfaced as a new therapeutic target. Hence, potent and selective NatD inhibitors would be valuable probes to interrogate its functions and therapeutical potential. However, there is no NatD inhibitor reported to date.

The co-crystal structure of the ternary NatD\CoA\SGRGK complex (PDB: 4U9W) revealed that its substrate peptide (SGRGK) is inserted into a highly acidic binding pocket of NatD, and biochemical studies supported the importance of the first 4 residues (SGRG) of the cognate substrate for specific recognition ^8^. Although the kinetic mechanism of NatD has not yet been elucidated, both NatA and NatE have been shown to follow an ordered Bi-Bi mechanism ^14–16^. Structural alignment of NatD with NatA infers a similar mechanism as NatA ^8^. Furthermore, upon mutation of an active-site C137, a previously suggested catalytic residue that forms an acetyl-NatD intermediate, hNatD remained largely active, supporting a direct transfer of the acetyl group through a Bi-Bi mechanism ^8^. Based on these observations, we hypothesized that bisubstrate analogues could provide potent inhibitors for NatD. Bisubstrate analogues have previously been used to help elucidate the catalytic mechanism of other NATs and to develop valuable tool compounds, usually displaying IC_50_ values within a low micromolar to high nanomolar range against *Sp*NatA ^14^, hNatB ^17^, hNatF ^18^, NatH ^19^. In the study presented here, we developed highly potent and selective NatD bisubstrate inhibitors. Moreover, the X-ray crystal structures of NatD bound to two bisubstrate inhibitors have been determined to elucidate the molecular interactions, thus paving the way for the structure-based development of more drug-like NatD inhibitors.

## RESULTS AND DISCUSSION

### Inhibitor Design

Bisubstrate analogues that covalently connect cofactor and short peptide substrate with a linker have been reported for targeting a variety of transferases, such as methyltransferases and acetyltransferases ^20,21^. Based on the co-crystal structure of the NatD/CoA/SGRGK ternary complex (PDB: 4U9W), the linear distance between the ⍺-nitrogen atom of the first Ser residue and the sulfur atom of CoA was measured to be about 3.1 Å (Figure 1) ^8^. Therefore, we proposed to connect the N⍺-amines of different peptide substrates and the thiol group of CoA through either an acetyl or propionyl linker to prepare bisubstrate analogues to mimic the transition state, which would be predicted to provide potent and selective inhibitors for NatD.

**Figure 1.**
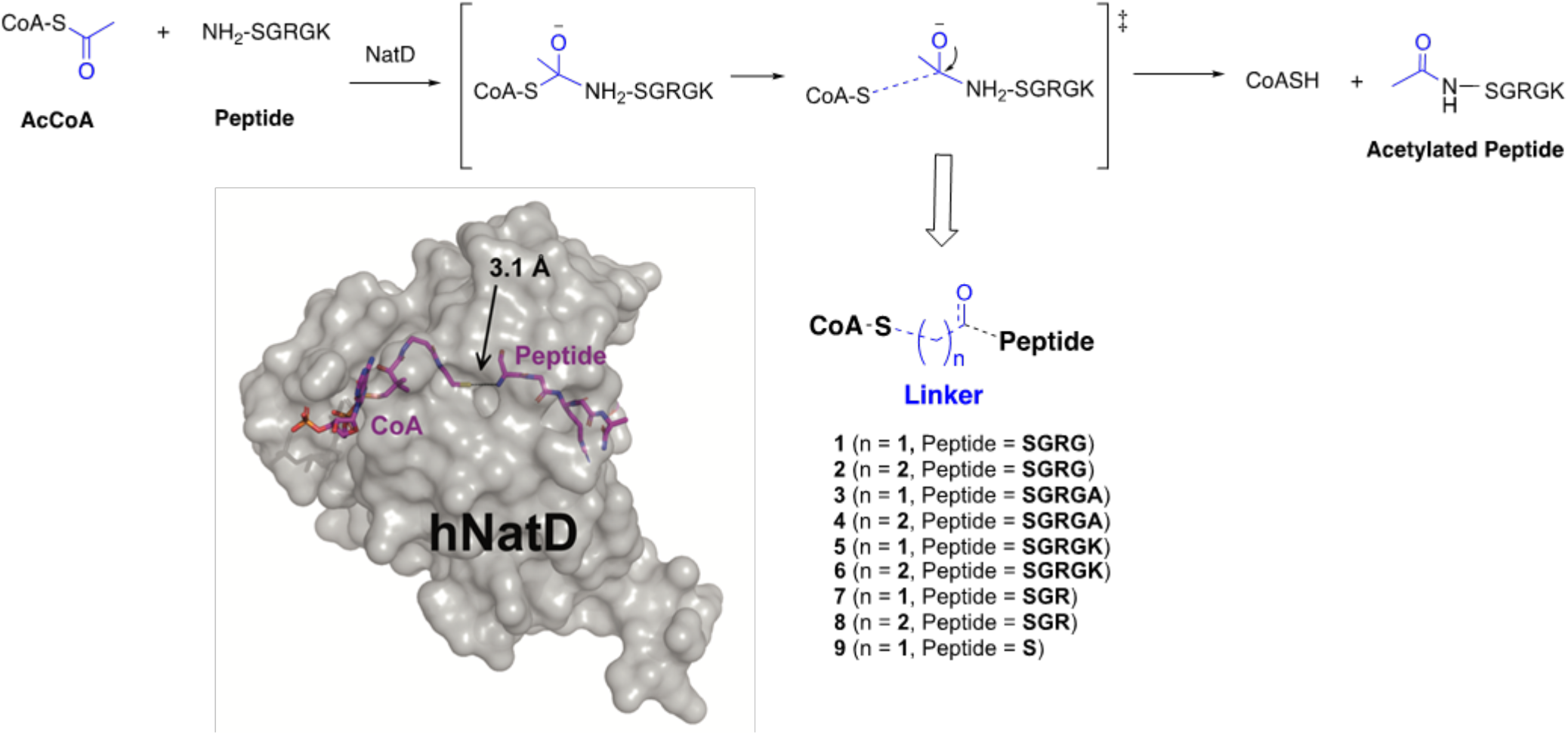
Inhibitor design strategy. The distance between the sulfur atom of CoA (magenta stick) and the Nα nitrogen atom of peptide (magenta stick) is 3.1 Å in the ternary complex of the NatD/CoA/SGRGK (PDB: 4U9W), where NatD is in surface view (grey). The designed NatD bisubstrate analogues **1**–**9** are illustrated.

### Inhibitor Synthesis

Peptides were prepared on Rink amide resin following a standard Fmoc solid-phase peptide synthesis protocol. 2-bromoacetic acid or 3-bromopropionic acid was coupled with the free N-terminal amine of the peptide on the resin. Subsequent cleavage with the cocktail consisting of trifluoroacetic acid (TFA)/water (H_2_O)/triisopropylsilane (95:2.5:2.5) and purification through high-performance liquid chromatography (HPLC) provided the purified bromopeptides, which were then reacted with coenzyme A trilithium salt dihydrate in the triethylammonium bicarbonate buffer ^20^. The resulting mixture was purified by HPLC to afford the desired bisubstrate analogues **1**–**9**.

**Scheme 1.**
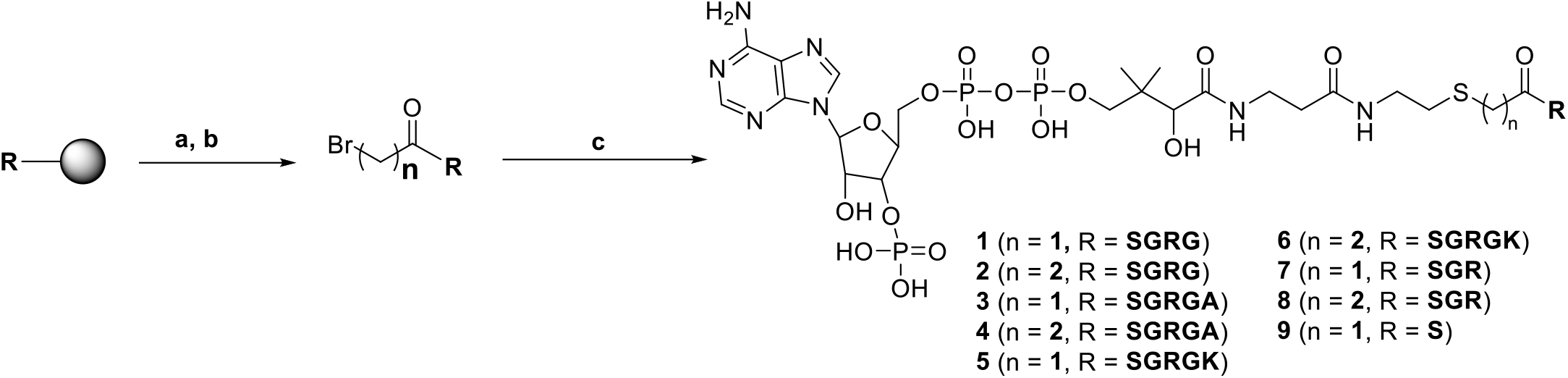
Synthetic route for the bisubstrate analogues. Reagents and conditions: (a) 2-Bromoacetic acid or 3-Bromopropionic acid, DIC, HOBt, DMF, r.t., overnight; (b) TFA/ TIPS/water(95/2.5/2.5), r.t., 4 h, 34–56% in two steps; (c) CoASH, triethylammonium bicarbonate buffer, pH 8.4 ± 0.1, r.t., 48 h, 33–55%.

### Structure-Activity Relationship Studies

All synthesized bisubstrate analogues were first evaluated with an established fluorescence assay ^22^. Initial testing was performed in the presence of both AcCoA and an H4-8 peptide (SGRGKGGK-CONH_2_) at their respective Michaelis constant (*K*_m_) values. But resulting IC_50_ values of compounds **1**–**8** were close to or lower than the concentration of NatD (45 nM) in the assay (Table 1, Figure 3Sa–b)), suggesting that they are slow and tight-binding inhibitors ^23, 24^. To understand the contribution of each component of the bisubstrate analogue to the NatD inhibition, we designed and prepared two control compounds **9** and **10**, where **9** connecting CoA with only Ser residue and **10** without the CoA moiety. Compound **9** displayed an IC_50_ value of 218 nM, which was over 4-fold reduction compared to those analogues containing 3–5 mer peptides. This result corroborated that the first three amino acids contribute significantly to the interaction with NatD as suggested by the ternary complex of NatD/CoA/SGRGK ^8^. Whereas, inhibitory activity of **10** lost over 1,000-fold even compared to **9**, supporting the importance of each moiety for potent inhibition of NatD. As fluorescence assay exhibited a narrow dynamic range (less than 2-fold) to fully characterize those slow and tight-binding inhibitors **1**-**8**, we applied an orthogonal radioactive assay to directly monitor the production of acetylated peptide under a similar condition with both AcCoA and H4-19 peptide substrate (SGRGKGGKGLGKGGAKRHR-COOH; GenScript) at their 4x*K*_m_ values as radioactive assay is generally more sensitive ^8^. The difference among **1**–**8** was more salient in the radioactive assay compared to the fluorescence assay, as tenfold difference was observed among IC_50_ values of **1**–**8** ranging from 4 nM to 41 nM for NatD (Table 1, Figure 3Sc,d). Inhibitory activity was reduced when only Ser was incorporated in the substrate moiety, displaying an IC_50_ value over 370 nM and supporting the importance of the first three amino acids. Meanwhile, removal of the CoA moiety also abolished the inhibitory activity, as propionyl-SGRGK **10** exhibited an IC_50_ of over 500 μM. Even under this saturated condition, **1**-**8** showed tight-binding inhibition as their IC_50_ values were below the concentration of NatD, Morrison’s quadratic equation was applied to analyze the concentration-response data to define their apparent *K*_i_ values ^24^. The most potent inhibitor **6** that links CoA and SGRGK peptide through a propionyl linker demonstrated a *K*_i,app_ value of 1.0 nM (Table 1, Figure 2a,b), which is 2-fold and 8.4-fold higher than **2** and **8**, respectively. For bisubstrate analogues containing either a tetrapeptide SGRG or pentapeptide SGRGK, a propionyl linker was more favorable than an acetyl linker as inhibitory activity was 2- to 4-fold better. However, acetyl linker was better than a propionyl linker for the bisubstrate ananlogs that only contained a tripeptide SGR, as **7** was 2-fold more potent than **8**. For the bisubstrate analogs containing a pentapeptide SGRGA, both linkers were acceptable and showed more similar potency. When three to five amino acids were incorporated in the bisubstrate analogue, both acetyl and propionyl linkers within the bisubstrate analogues can be accommodated by NatD.

**Table 1.** Inhibition activities of the bisubstrate inhibitors. Note: a) Inhibition activities were measured in duplicate (n = 2) in the fluorescence assay with both AcCoA and peptide substrate were at their respective *K*_m_ values. b) IC_50_ values were determined in triplicate (n = 3) in the radioactive assay with both AcCoA and peptide substrate were at their respective 4x*K*_m_ values. C) Apparent *K*_i_ values were analyzed by fitting concentration–response data obtained in the radioactive assay to Morrison’s quadratic equation for **1**-**8**. IC_50_ and *K*_i,app_ values were presented as mean ± standard deviation (SD). NA: not applied.

| ID | Structure | IC<sub>50</sub> (nM)<sup>a</sup> | IC<sub>50</sub> (nM)<sup>b</sup> | Morrison $K_{i,app}$ (nM)<sup>c</sup> |
| --- | --- | --- | --- | --- |
| **1** | CoA-C2-SGRG | 48 ± 2.5 | 27 ± 3.8 | 4.4 ± 0.85 |
| **2** | CoA-C3-SGRG | 48 ± 8.4 | 7.1 ± 1.8 | 2.1 ± 0.32 |
| **3** | CoA-C2-SGRGA | 41 ± 3.8 | 25 ± 3.4 | 6.2 ± 1.5 |
| **4** | CoA-C3-SGRGA | 32 ± 2.5 | 35 ± 5.4 | 8.7 ± 1.8 |
| **5** | CoA-C2-SGRGK | 35 ± 6.0 | 20 ± 3.2 | 4.4 ± 0.58 |
| **6** | CoA-C3-SGRGK | 40 ± 7.7 | 4.1 ± 0.40 | 1.0 ± 0.11 |
| **7** | CoA-C2-SGR | 29 ± 5.6 | 16 ± 4.9 | 3.7 ± 0.71 |
| **8** | CoA-C3-SGR | 40 ± 3.9 | 41 ± 6.7 | 8.4 ± 1.6 |
| **9** | CoA-C2-S | 218 ± 28 | > 370 | NA |
| **10** | Propionyl-SGRGK | 330 ± 35 μM | > 500 μM | NA |

**Figure 2.**
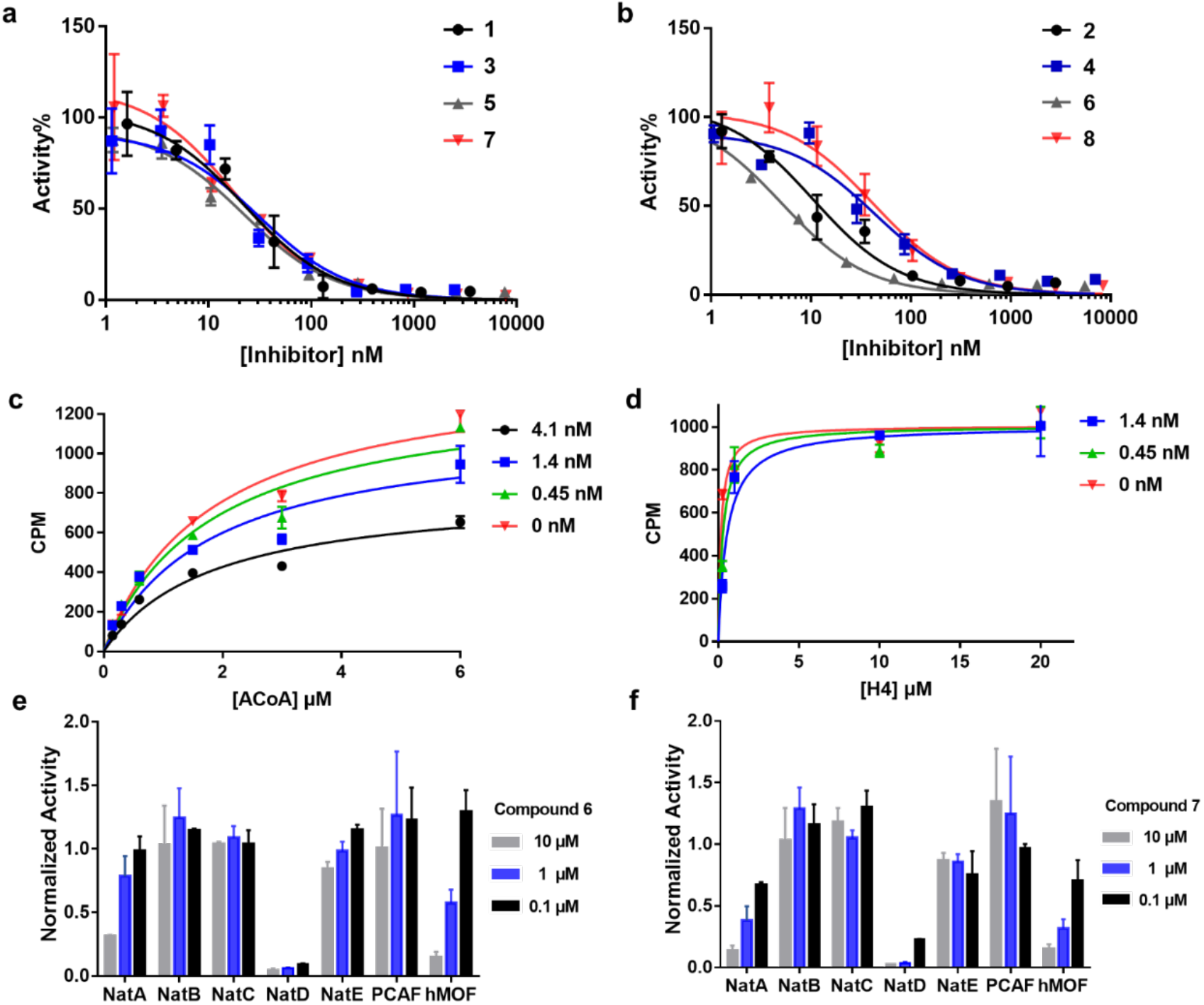
Biochemical characterization of bisubstrate analogues via the radioactive assay. a) Concentration-response plots of bisubstrate analogues with an acetyl linker under 4x*K*_m_ of H4-19 peptide and AcCoA condition fitted to Morrison’s quadratic equation (n=3). b) Concentration-response plots of bisubstrate analogues with a propionyl linker under 4x*K*_m_ of H4-19 peptide and AcCoA condition (n=3). c) CoA-C3-SGRGK (**6)** is noncompetitive with respect to AcCoA. d) **6** is competitive with respect to H4. e,f) Selectivity study **6** and **7** against a panel of protein acetyltransferases in triplicate (n=3).

**Figure 3.**
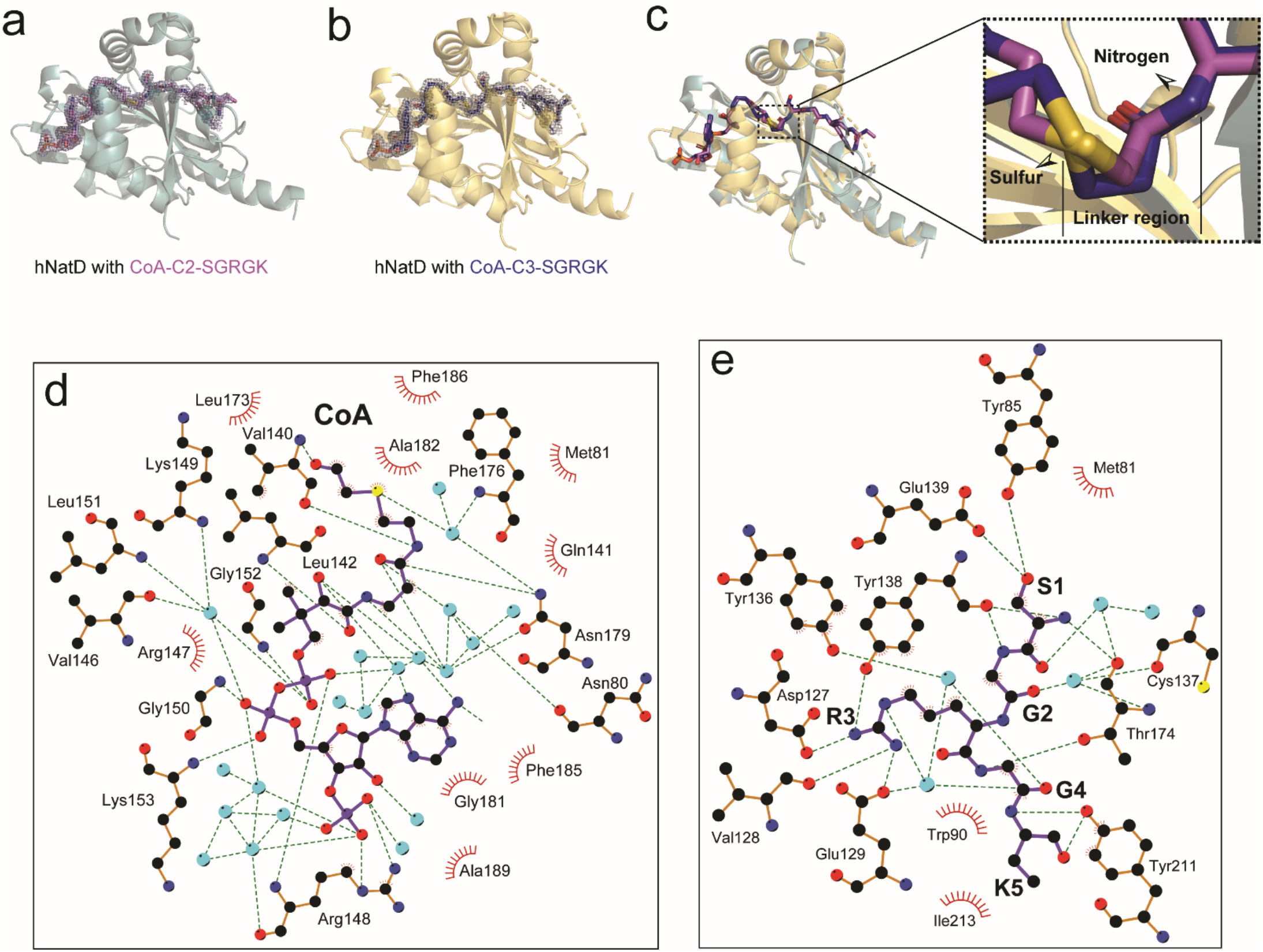
Structures of hNatD with Inhibitors. (a) Structure of hNatD with **5** (CoA-C2-SGRGK) bound. hNatD is shown in cyan as a cartoon and CoA-C2-SGRGK as magenta sticks. The 2mF_obs_-DF_cal_ electron density around the bisubstrate is shown at a contour level of 1σ (gray mesh). (b) Structure of hNatD with **6** (CoA-C3-SGRGK) bound. hNatD is shown in yellow as a cartoon and CoA-C3-SGRGK as blue sticks. The 2mF_obs_-DF_cal_ electron density around the bisubstrate is shown at a contour level of 1σ (gray mesh). (c) Overlay of the structure in (a) and (b). The zoom-in view shows the positions of the sulfur atom of CoA and the nitrogen of the peptide amino group. (d) Interaction between CoA and hNatD residues is generated with LIGPLOT Hydrogen bonds are indicated by dashed green lines, and van der Waals interactions are indicated with red semicircles. Waters molecules are shown as cyan spheres. (e) Interaction between SGRGK and hNatD residues is generated with LIGPLOT. Hydrogen bonds are indicated by dashed green lines, and van der Waals interactions are indicated with red semicircles. Waters molecules are shown as cyan spheres.

### Inhibition mechanism

The most potent compound **6** was selected to investigate the inhibition mechanism. As shown in figure 2c, compound **6** exhibited a noncompetitive inhibition pattern with AcCoA. The data were globally fit to noncompetitive inhibition equation as a preferred model, producing the best fit parameters, α*K*_I_ = 2.1 ± 0.38 nM and *K*_m_ = 0.64 ± 0.07 nM. In contrast, compound **6** exhibited a competitive inhibition pattern with peptide substrate (Figure 2d). The data were globally fit to competitive inhibition equation as a preferred model, producing the best fit parameters, *K*_i_ = 0.61 ± 0.22 nM and *K*_m_ = 0.16 ± 0.03 nM. For a two substrate enzyme with an ordered binding mechanism where substrate A preferentially binds before substrate B, it is expected that a bisubstrate analog will be competitive with substrate A and noncompetitive with substrate B ^25^. Thus, our results suggested NatD undergoes an ordered Bi-Bi mechanism, in which the peptide substrate preferentially binds NatD first, followed by AcCoA.

### Selectivity Study

Two of the most potent bisubstrate analogues **6** and **7** were chosen to examine their selectivity against a panel of closely-related N-terminal acetyltransferases, including NatA, NatB, NatC, NatE; and protein lysine acetyltransferases PCAF and hMOF, which all harbor an Ac-CoA binding motif. Both **6** and **7** did not show any inhibition towards NatB, NatC, NatE, and PCAF at 10 μM (Figure 2e,f), indicating over 10,000-fold selectivity for NatD. Compound **6** demonstrated 10–40% of inhibition against both NatA and hMOF at 1.0 μM, indicating over 1,000 fold selectivity for NatD over NatA and hMOF. **7** displayed IC_50_ values over 0.1 μM against both NatA and hMOF, suggesting that it was less selective than **6**. As NatA is a founding member of the NAT family that prefers to acetylate substrates starting with Ser or Ala, decreased selectivity of **7** compared to **6** is informative in the future development of selective NatD inhibitors.

### Co-crystal Structures of 5 and 6 with NatD

To understand the molecular interactions between the bisubstrate inhibitors and NatD, the X-ray co-crystal structures of NatD complexed with **5** (PDB ID: 7KD7) and **6** (PDB ID: 7KPU) were obtained at 1.44 Å and 1.43 Å resolution, respectively **(Figure 3a-b)**. The structure refinement statistics for these structures can be found in **Table S1**. Overall, both structures showed high similarity to the previously reported NatD/CoA/SGRGK ternary complex (PDB ID: 4U9W) ^8^. Structural alignment of the **5-NatD** and **6-NatD** binary complexes with the NatD/CoA/SGRGK ternary complex gave RMSD values of 0.255 Å and 0.217 Å, respectively. Interestingly, despite the different length of the linker region within the two bisubstrate inhibitors, the sulfur atom of the CoA and the nitrogen atom of the peptide Nα-amino group remain in the same position (**Figure 3c)**. Thus, such similar positioning of these two atoms explains the comparable potency of the inhibitors with two different linkers as described above. Also noteworthy is that CoA-C2-SGRGK and CoA-C3-SGRGK adopt slightly different CoA binding modes, especially with respect to the β-mercaptoethylamine group **(Figure 3c)**. Thus, we speculate that the flexibility of this CoA binding region endows NatD the ability to accept a bisubstrate inhibitor CoA-C3-SGRGK with a relatively long linker. The structures also reveal why **9** and **10** poorly inhibit NatD, since both the peptide and CoA portion of the inhibitor make important contributions to inhibitor binding. The tight binding of bisubstrate analogues further supports the Bi-Bi mechanism of NatD.

Given the similarity between both NatD/inhibitor structures, we focused on the structure of NatD with CoA-C2-Ser^1^-Gly^2^-Arg^3^-Gly^4^-Lys^5^ to further examine the molecular basis of NatD inhibition. An extensive H-bonding network is observed for CoA recognition involving several water molecules and NatD residues Asn80, Val140, Leu142, Val146, Arg148, Lys149, Gly150, Leu151, Gly152, Lys153, Phe176, and Asn179 **(Figure 3d)**. Hydrophobic interactions are also observed from NatD residues Met81, Gln141, Arg 147, Leu173, Gly181, Ala182, Phe185, Phe186, and Ala189. These extensive interactions explain why inhibitory activity drops significantly when the CoA portion is removed (**Table 1**, compare **6** and **10**). NatD interacts with the peptide portion of the bisubstrate inhibitor through hydrogen bonding to the backbone of residues 1– 5, and the sidechains of Ser^1^ and Arg^3^. These interactions involve several water molecules and NatD residues Tyr85, Asp127, Val128, Glu129, Tyr136, Cys137, Tyr138, Gln139, Thr174, and Tyr121 **(Figure 3e)**. In contrast, van der Waals interactions are significantly more limited, involving only several residues, including Met80, Trp90, Ile213. The extensive NatD interaction with the first three residues is consistent with the biochemical inhibitory results: inhibition potency depends on the inclusion of Ser^1^ through Arg^3^ (**Table 1**, compare **9** to **1–8**). The NatD/inhibitor structures also reveal that the sidechain of Lys^5^ does not mediate any interactions with the enzyme, which is consistent with the biochemical findings that a minimal effect on inhibitor potency was observed when Lys^5^ was replaced with Ala (**Table 1**, compare **3–4** to **5–6**). Taking the structures together with the binding data, we propose that compounds that mimic at least the mercapto group of acetyl-CoA and the SGR sequence of the peptide with some toleration for flexibility linking these regions could represent suitable drug-like lead molecules for NatD inhibition.

### Cellular Target Engagement

Although the inherent properties of bisubstrate inhibitors preclude their direct use in cellular studies as they are not be able to directly penetrate the cell membrane, we took advantage of digitonin permeabilization to promote cell uptake of compounds **2** and **6** for a preliminary investigation on cellular target engagement ^26^. At 100 μM of compounds **2** and **6** with 2 h incubation, we were able to introduce the compounds into digitonin (5 μg/mL) treated HCT116 cells (Figure S4) ^27^. Although the amount of **2** and **6** that could enter cells could not be determined, both **2** and **6** induced a higher thermal stabilization (Δ*T*_agg_ = +1.3 °C and +1.7 °C, respectively) using a cellular thermal shift assay (CETSA) compared to the control (Figure 4) ^28,29^. This result demonstrated that **2** and **6** could engage NatD in cells. Future efforts will focus on the introduction of cell-permeable peptides at the C-terminal end of the bisubstrate inhibitors to improve the cell penetration, together with a comprehensive SAR of **6** for future optimization.

**Figure 4.**
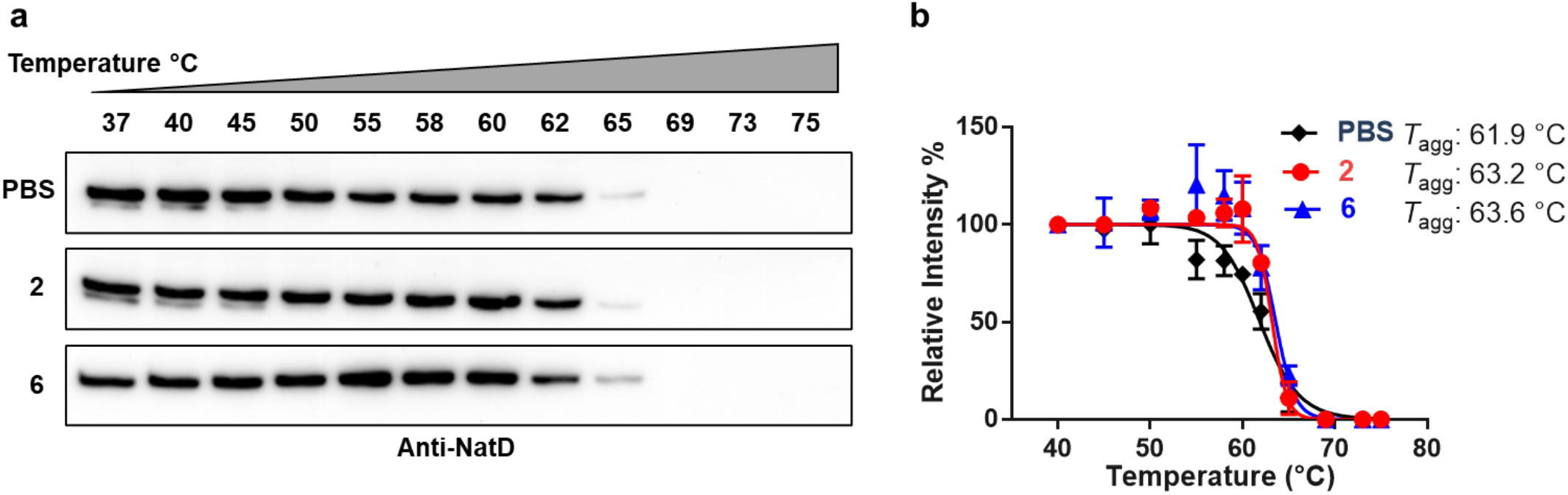
Cellular thermal shift assay. a) Thermal stability of endogenous NatD by immunoblot using anti-NatD antibody (Invitrogen PIPA520882) when treated with **2** or **6** or PBS buffer for 2 h at 100 μM after digitonin (5 μg/mL) permeabilization for 20 min. b) Quantified the relative NatD band intensities of the immunoblot in duplicates.

## CONCLUSION

In this work, we designed and synthesized the first series of potent and selective NatD bisubstrate inhibitors **1–8**, exhibiting *K*_i,app_ values ranging from 1.0–8.7 nM in the radioactive assay. The most potent inhibitor CoA-C3-SGRGK showed a *K*_i,app_ of 1.0 ± 0.11 nM, exhibiting about 1,000-fold selectivity for NatD over other acetyltransferases, including NatA-C, NatE, hMOF, and PCAF. Compared to reported bisubstrate inhibitors for other NATs, including NatA and NatE using a similar bisubstrate strategy ^20^, the high potency of NatD bisubstrate inhibitors further establish the uniqueness of NatD and strengthen the possibility to develop potent and specific inhibitors for NatD. Furthermore, the co-crystal structures of NatD in complex with CoA-C3-SGRGK and CoA-C2-SGR clearly demonstrate that the bisubstrate inhibitors engaged both substrate and cofactor AcCoA binding sites. The structural observations reveal that the NatD active site is specifically tailored for its histone substrate, explaining the selectivity of the NatD bisubstrate inhibitors. Comparable inhibitory activities and minor differences in the AcCoA binding site for bisubstrate analogues with either an acetyl or propionyl linker suggest active site plasticity, which may result from a subtle conformational change within the cofactor binding site of NatD. Moreover, these co-crystal structures of NatD-inhibitor binary complexes provide a structural foundation to guide the future development of drug-like NatD inhibitors.

## EXPERIMENTAL SECTION

### Chemistry General Procedures

All chemicals and solvents were purchased from commercial suppliers and used without further purification unless stated otherwise. Preparative high pressure liquid chromatography (RP-HPLC) was performed on an Agilent 1260 Series system. Systems were run with 0–95% methanol/water gradient with a 0.1% TFA modifier. High-resolution Matrix-assisted laser desorption/ionization (MALDI) spectra were performed on a 4800 MALDI TOF/TOF mass spectrometry (Sciex) at the Mass Spectrometry and Purdue Proteomics Facility (PPF), Purdue University. Peptides were synthesized on a CEM Liberty Blue peptide synthesizer. Compounds were also characterized and confirmed by TLC-MS or MALDI-MS. The purity of final compounds was confirmed on a Waters LC-MS system and/or Agilent 1260 Series system. Systems were run with 0–40% methanol/water gradient with 0.1% TFA. The purity of all target compounds showed >95%.

### General procedure A for solid-phase peptide synthesis

The Peptides were synthesized using a Liberty Blue automated microwave peptide synthesizer (CEM Corp., Matthews, NC, USA) following a standard Fmoc protocol ^30^. A Rink Amide MBHA resin (0.05 mmol) was used as solid support and placed in the microwave tube. Standard couplings of amino acids were carried out at 0.2 M in DMF and external amino acids at 0.1 M in DMF using 0.5 M DIC and 1.0 M Oxyma in DMF for activation and 20% piperidine in DMF for deprotection. The resin was transferred to a filter-equipped syringe, washed with CH_2_Cl_2_ (3 mL) and MeOH (3 mL) three times, and dried under air.

### General procedure B for the synthesis of bromoacetylated peptide.^20^

To a suspension of peptide on resin (0.05 mmol, 1.0 equiv) in DMF (2 mL) were added corresponding acid (0.1 mmol, 2 equiv), DIC (15.5 μL, 0.1 mmol, 2 equiv), and HOBt (13.5 mg, 0.1 mmol, 2 equiv). The suspension was shaken at room temperature for 20 h. The solvent was filtered, and the resulting resin was washed with CH_2_Cl_2_ (3 mL) and MeOH (3 mL) three times and dried under air. The dried peptide was cleaved from the resin using a cleavage cocktail (TFA/TIPS/ddH2O = 95/2.5/2.5 v/v) (5 mL) for 0.05 mmol scale of resin for 4 h. The suspension was filtered, washed with TFA (2 mL) and the volatiles of the filtrate was removed under N_2_. The peptide solution was precipitated with cold anhydrous ether (10 mL), centrifuged at 4200 rpm for 10 min. The supernatant was discarded, and the pellet was washed with cold ether, centrifuged, removed supernatant, and air-dried. The dried peptide was dissolved in ddH2O (5 mL) and filtered through a 0.2 μM filter membrane. The filtered sample solution was purified by preparative reversed-phase high performance liquid chromatography (RP-HPLC) using an Agilent 1260 Series system with a C18 column (5 μm, 10 mm × 250 mm) at a flow rate of 4.0 mL/min. Two mobile phases (mobile phase A consisting of 0.1% trifluoroacetic acid in ddH2O; mobile phase B consisting of methanol) were used and monitored at 214 and 256 nm. An injection volume of 400 μL of the solution was injected into the column. The desired fractions were evaporated, and the resulting solution was frozen (–85 °C) and lyophilized.

### General procedure C for synthesis of the bisubstrate analogues^20^

The bromoacetylated peptide (1 equiv) was dissolved in 1 mL triethylammonium bicarbonate buffer, pH 8.4 ± 0.1. The mixture was added with coenzyme A trilithium salt dihydrate (2 equiv), and the solution was allowed to react for 4 h at room temperature and then overnight at 4 °C. 100 uL of the reaction solution was purified by HPLC to obtain the desired product, which was detected by MALDI.

Compound **1** (5.1 mg, 43% yield) was synthesized by following the general procedure C as a foamy white solid. MALDI-TOF (positive) *m*/*z*: calcd for C_36_H_63_N_15_O_22_P_3_S^+^ [M + H]^+^ *m*/*z* 1182.3200, found *m*/*z* 1182.3802. Compound **2** (4.3 mg, 36 % yield) was synthesized by following the general procedure C as a foamy white solid. MALDI-TOF (positive) *m*/*z*: calcd for C_37_H_65_N_15_O_22_P_3_S^+^ [M + H]^+^ *m*/*z* 1196.3357, found *m*/*z* 1196.3724. Compound **3** (5.6 mg, 45% yield) was synthesized by following the general procedure C as a foamy white solid. MALDI-TOF (positive) *m*/*z*: calcd for C_39_H_68_N_16_O_23_P_3_S^+^ [M + H]^+^ *m*/*z* 1253.3571, found *m*/*z* 1253.3676. Compound **4** (4.8 mg, 38% yield) was synthesized by following the general procedure C as a foamy white solid. MALDI-TOF (positive) *m*/*z*: calcd for C_40_H_70_N_16_O_23_P_3_S^+^ [M + H]^+^ *m*/*z* 1267.3728, found *m*/*z* 1267.4188. Compound **5** (6.2 mg, 47% yield) was synthesized by following the general procedure C as a foamy white solid. MALDI-TOF (positive) *m*/*z*: calcd for C_42_H_75_N_17_O_23_P_3_S^+^ [M + H]^+^ *m*/*z* 1310.4150, found *m*/*z* 1310.4417. Compound **6** (4.8 mg, 36% yield) was synthesized by following the general procedure C as a foamy white solid. MALDI-TOF (positive) *m*/*z*: calcd for C_43_H_77_N_17_O_23_P_3_S^+^ [M + H]^+^ *m*/*z* 1324.4306, found *m*/*z* 1324.4221. Compound **7** (4.7 mg, 42% yield) was synthesized by following the general procedure C as a foamy white solid. MALDI-TOF (positive) *m*/*z*: calcd for C_34_H_60_N_14_O_23_P_3_S^+^ [M + H]^+^ *m*/*z* 1125.2986, found *m*/*z* 1125.2793. Compound **8** (3.8 mg, 33% yield) was synthesized by following the general procedure C as a foamy white solid. MALDI-TOF (positive) *m*/*z*: calcd for C_35_H_62_N_14_O_23_P_3_S^+^ [M + H]^+^ *m*/*z* 1139.3142, found *m*/*z* 1139.2950. Compound **9** (4.7 mg, 51% yield) was synthesized by following the general procedure C as a foamy white solid. MALDI-TOF (positive) *m*/*z*: calcd for C_26_H_45_N_9_O_19_P_3_S^+^ [M + H]^+^ *m*/*z* 912.1760, found *m*/*z* 912.1606. Compound **10** (3.1 mg, 55% yield) was synthesized by following the general procedure B as a foamy white solid. MALDI-TOF (positive) *m*/*z*: calcd for C_22_H_43_N_10_O_7_^+^ [M + H]^+^ *m*/*z* 559.3311, found *m*/*z* 559.3316.

### Protein Expression and Purification

Expression and purification of human PCAF, hMOF, NatA-E except for NatC, were performed as previously described ^8,17,31–34^.

#### NatC

Ternary *S. pombe* NatC encoding NAA30^FL^, NAA35^31-708,^ and NAA38^48-116^, was expressed in *E. coli* cells, and purified as follows. Cells were harvested by centrifugation, resuspended, and lysed by sonication in lysis buffer containing 25 mM Tris, pH 8.0, 300 mM NaCl, 10 mg/mL PMSF (phenylmethanesulfonylfluoride). The lysate was clarified by centrifugation and passed over a nickel resin (Thermo Scientific), which was subsequently washed with ~10 column volumes of wash buffer containing 25 mM Tris, pH 8.0, 300 mM NaCl, 20 mM imidazole, 10 mM 2-mercaptoethanol. The protein was eluted with buffer containing 25 mM Tris, pH 8.0, 300 mM NaCl, 200 mM imidazole, 10 mM βME. After elution, His-tagged Ulp1 protease was added to the eluent to cleave the SUMO tag. The eluent was further dialyzed into a buffer containing 25 mM sodium citrate monobasic, pH 5.5, 10 mM NaCl and 10 mM 2-mercaptoethanol. Protein was purified with a 5-mL HiTrap SP ion-exchange column and eluted with a salt gradient (10–1000 mM NaCl). Peak fractions were concentrated to ~ 0.5 mL with a 50 kDa concentrator (Amicon Ultra, Millipore), and loaded onto an S200 gel-filtration column (GE Healthcare) in a buffer containing 25 mM HEPES, pH 7.0, 200 mM NaCl, 1 mM dithiothreitol (DTT). Peak fractions were concentrated to ~ 15 mg/mL as measured by UV280 and flash-frozen for storage at −80 °C until use.

### Fluorescence Assays

A fluorescence-based assay was applied to study the IC_50_ values for all the compounds. The assay was performed under the following conditions in a final well volume of 40 μL: 25 mM HEPES (pH = 7.5), 150 mM NaCl, 0.01% Triton X-100, 0.05 μM NatD, 0.5 μM AcCoA, and 15 μM ThioGlo4. The inhibitors were added at concentrations ranging from 0.15 nM to 10 μM. After 30 min incubation, reactions were initiated by the addition of 5.0 μM H4-8 peptide. Fluorescence was monitored on a BMG ClariOtar microplate reader with excitation 400 nm and emission 465 nm. Data were processed by using GraphPad Prism software 7.0.

### Radioisotopic Acetyltransferase Assay

FL NatD was used for activity assays. NatD Acetyltransferase assays were carried out in 25 mM HEPES, pH 7.5, 150 mM NaCl, and 1 mM DTT. The H4 substrate peptide used in the assay corresponds to the first 19 residues of human H4 (NH_2_-SGRGKGGKGLGKGGAKRHR-COOH; GenScript). In the assay, 50 nM of hNatD was mixed with 2 μM of radiolabeled [^14^C]acetyl-CoA (4 mCi/mmol; PerkinElmer Life Sciences), 20 μM of the peptide, and inhibitors of concentrations ranging from 0.15 nM to 10 μM, for a reaction of 30 minutes at room temperature. To quench the reaction, the reaction solution was applied onto negatively charged P81 paper disks (SVI, St vincent’s institute medical research) to trap the peptides, and the paper disks were immediately placed in wash buffer (10 mM HEPES, pH 7.5). The paper disks were washed three times, at 5 minutes per wash, to remove unreacted acetyl-CoA. The papers were then dried with acetone and added to 4 mL of scintillation fluid, and the signal was measured with a Packard Tri-Carb 1500 liquid scintillation analyzer. Each reaction was performed in triplicate. IC_50_ statistic values were determined with GraphPad Prism software 7.0 ^8^.

### Inhibition Mechanism Study

The experiment procedures are the same as described for the radioactive assay. In the AcCoA concentration fixed assay, 50 nM of hNatD was mixed with 2 μM of radiolabeled [^14^C]acetyl-CoA, the H4 peptide (0.25, 1.0, 10 and 20 μM), and inhibitors (0, 0.45 and 1.4 nM), for a reaction of 30 minutes at room temperature. In the H4 peptide concentration fixed assay, 50 nM of hNatD was mixed with 20 μM of H4 peptide, the [^14^C]acetyl-CoA (0.15, 0.3, 0.6, 1.5, 3 and 6 μM), and inhibitors (0, 0.45, 1.4 and 4.1 nM), for a reaction of 13 minutes at room temperature.

### Selectivity Assays

The selectivity studies of NatA, NatB, NatC, NatE, PCAF, and hMOF were performed as follows. 100 nM of hNatA was mixed with 30 μM of [^14^C]acetyl-CoA and 30 μM of either H4 peptide or SASE peptide (NH_2_-SASEAGVRWGRPVGRRRRP-COOH; GenScript), with none, 0.1 μM, 1 μM or 10 μM of inhibitors, for a 12-minute reaction at room temperature, in the buffer containing 75 mM HEPES, pH 7.5, 120 mM NaCl, 1mM DTT.

100 nM of hNatB was mixed with 50 μM [^14^C]acetyl-CoA and 50 μM of MDVF peptide (NH_2_-MDVFMKGRWGRPVGRRRRP-COOH, GenScript), with none, 0.1 μM, 1 μM or 10 μM of inhibitors, for a 10-minute reaction at room temperature, in the buffer containing 75 mM HEPES, pH 7.5, 120 mM NaCl, 1 mM DTT.

50 nM of *Sp*NatC was mixed with 30 μM [^14^C]acetyl-CoA and 10 μM of MLRF peptide (NH_2_-ML RFVTKRWGRPVGRRRRPCOOH, GenScript), with none, 0.1 μM, 1 μM or 10 μM of inhibitors, for a 5-minute reaction at room temperature, in the buffer containing 75 mM HEPES, pH 7.0, 120 mM NaCl, 1 mM DTT.

300 nM of hNatE was mixed with 50 μM [^14^C]acetyl-CoA and 100 μM of MLGP peptide (NH_2_-MLGPEGGRWGRPVGRRRRP-COOH, GenScript), with none, 0.1 μM, 1 μM or 10 μM of inhibitors, for a 40-minute reaction at room temperature, in the buffer containing 75 mM HEPES, pH 7.0, 120 mM NaCl, 1 mM DTT.

100 nM of PCAF was mixed with 50 μM [^14^C]acetyl-CoA and 400 μM of H4 peptide, with none, 0.1 μM, 1 μM or 10 μM of inhibitors, for a 20-min reaction at room temperature, in the buffer containing 40 mM Tris, pH 8.0, 100 mM NaCl, 1 mM DTT and 2 mg/mL BSA.

50 nM of hMOF was mixed with 50 μM [^14^C]acetyl-CoA and 400 μM of H4 peptide, with none, 0.1 μM, 1 μM or 10 μM of inhibitors, for a 20-min reaction at room temperature, in the buffer containing 100 mM Tris, pH 8.0, 50 mM NaCl, 1 mM DTT, 800 μM cysteine and 0.25 mg/mL BSA.

### Co-crystallization and Structure Determination

For co-crystallization and structure determination of **5** and **6** with NatD, a truncation construct of hNatD^17–220^ was used and purified similarly as described above. 10 mg/mL of purified hNatD^17-220^ was incubated with 1 mM of either **5** or **6** for 30 minutes in ice before the crystal setup. The best crystals of hNatD^17-220^ with **5** bound was obtained with hanging-drop vapor diffusion at 20 °C in a well containing 0.1 M Bis-Tris, pH 5.5, 2 M Ammonium sulfate, in a drop containing a 1:1.5 mixture of protein to a well solution. The best crystals of hNatD^17-220^ with **6** bound was obtained with hanging-drop vapor diffusion at 20 °C in a well solution containing 0.1 M Bis-Tris, pH 5.5, 2 M Ammonium sulfate, in a drop containing a 1.25:1 mixture of protein to a well solution. All crystals were cryoprotected by transferring them to their respective well solutions supplemented with 20% glycerol before being flash frozen in liquid nitrogen. Data were collected at the Advanced Photon Source (beamline 24-ID-E) and processed using HKL2000 ^35^.

### Structure Determination and Refinement

Both structures were determined by molecular replacement using the structure of NatD/CoA/SGRGK (PDB: 4U9W) with ligands and solvent molecules removed from the search model. Molecular replacement was done using Phaser in Phenix ^36^. Initial Manual model building was done in Coot ^37^ and all subsequent rounds of refinement were performed using Phenix refine and Coot interchangeably. Refinement statistics can be found in **Table S1.** The final model and structure factors were submitted to the Protein Data Bank. Distance calculations, as well as three-dimensional alignment r.m.s. deviations and graphics were generated in PyMOL (http://www.pymol.org/).

### Cellular thermal shift assay

The assay was performed based on a modified protocol for the cell lysate CETSA experiments ^28^. In brief, cultured HCT116 cells (5.0 × 10^7^) was incubated with 5 μg/mL digitonin at 37 °C for 20 min ^26^. The medium was aspirated and replenished with fresh medium with PBS or Inhibitors at 100 μM. After 2 h, the cells were harvested and washed with PBS and resuspended in 1 mL of PBS. 300 μL of the cells solution were directly lysed by three freeze-thaw cycles with liquid nitrogen and clarified by centrifugation at 20,000g for 20 min at 4 °C. The supernatant was analyzed with MALDI-MS. Aliquots of 50 μL from each condition were distributed into PCR strip tubes and heated at 37, 40, 45, 50, 55, 58, 60, 62, 65, 69, 73, and 75 °C for 3 min before being cooled to room temperature for 3 min. The cells were lysed by three freeze-thaw cycles with liquid nitrogen and clarified by centrifugation at 20,000g for 20 min at 4 °C. The soluble fractions (lysate) were then analyzed by immunoblotting using anti-NatD antibody (Invitrogen PIPA520882) and quantified using ImageJ ^29^.

## ASSOCIATED CONTENT

### Supporting Information

The Supporting Information is available free of charge on the ACS Publications website.

Figure S1-2. and HPLC spectra of compounds **1**-**10**; Figure S3. IC_50_ fitting curves of compounds; Figure S4. Cell permeability study; Table S1. Data statistics for NatD crystal structures in complex with inhibitors. Molecular formula strings for all synthesized inhibitors.

### Accession Codes

The coordinates for the structure of human NatD in complex with compound **5 (**PDB ID: 7KD7**) and 6 (**PDB ID: 7KPU) have been deposited in the Protein Data Bank. Authors will release the atomic coordinates and experimental data upon article publication.

### Author Information

Corresponding Author

*Phone: (765) 494 3426.

### Author Contributions

Y.D. synthesized and characterized all compounds described in the manuscript. Y.H. and Y.D. performed fluorescence assay. Z.H. contributed to the synthesis. Y.D. and S.D. prepared manuscript figures and text. S.D. performed the radioactivity assay and obtained the co-crystal structures. S.D. and S.M.G. performed the selectivity study. R.M. designed and supervised experiments by S.D. and S.M.G. and prepared manuscript text. R.H. developed the concept, designed and supervised experiments by Y.D., Y.H., and Z.H., and prepared manuscript text and figures. All authors have read and approved the final version of the manuscript.

#### Notes

The authors declare no competing financial interest.

## ACKNOWLEDGMENT

The authors acknowledge the support from NIH grant R35GM118090 (RM) and proteomics core facility at Purdue University Center for Cancer Research (P30 CA023168). We also thank the support from the Department of Medicinal Chemistry and Molecular Pharmacology (RH) and Leah Gottlieb for her assistance in obtaining proteins of PCAF and hMOF for the inhibitor selectivity studies.

## ABBREVIATIONS

NatA: protein N-terminal acetyltransferase A
CoA: coenzyme A
AcCoA: acetyl coenzyme A
rt: room temperature
TFA: trifluoroacetic acid.

---

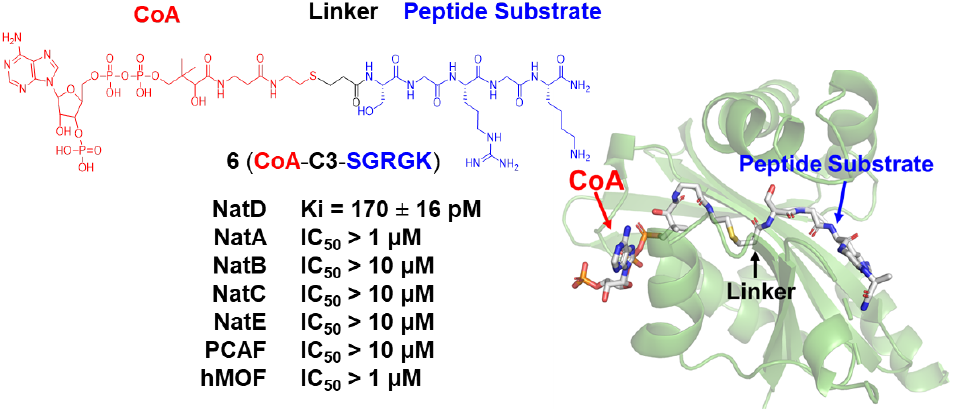
Table of Contents graphic

## Supporting Information

**Figure S1.**
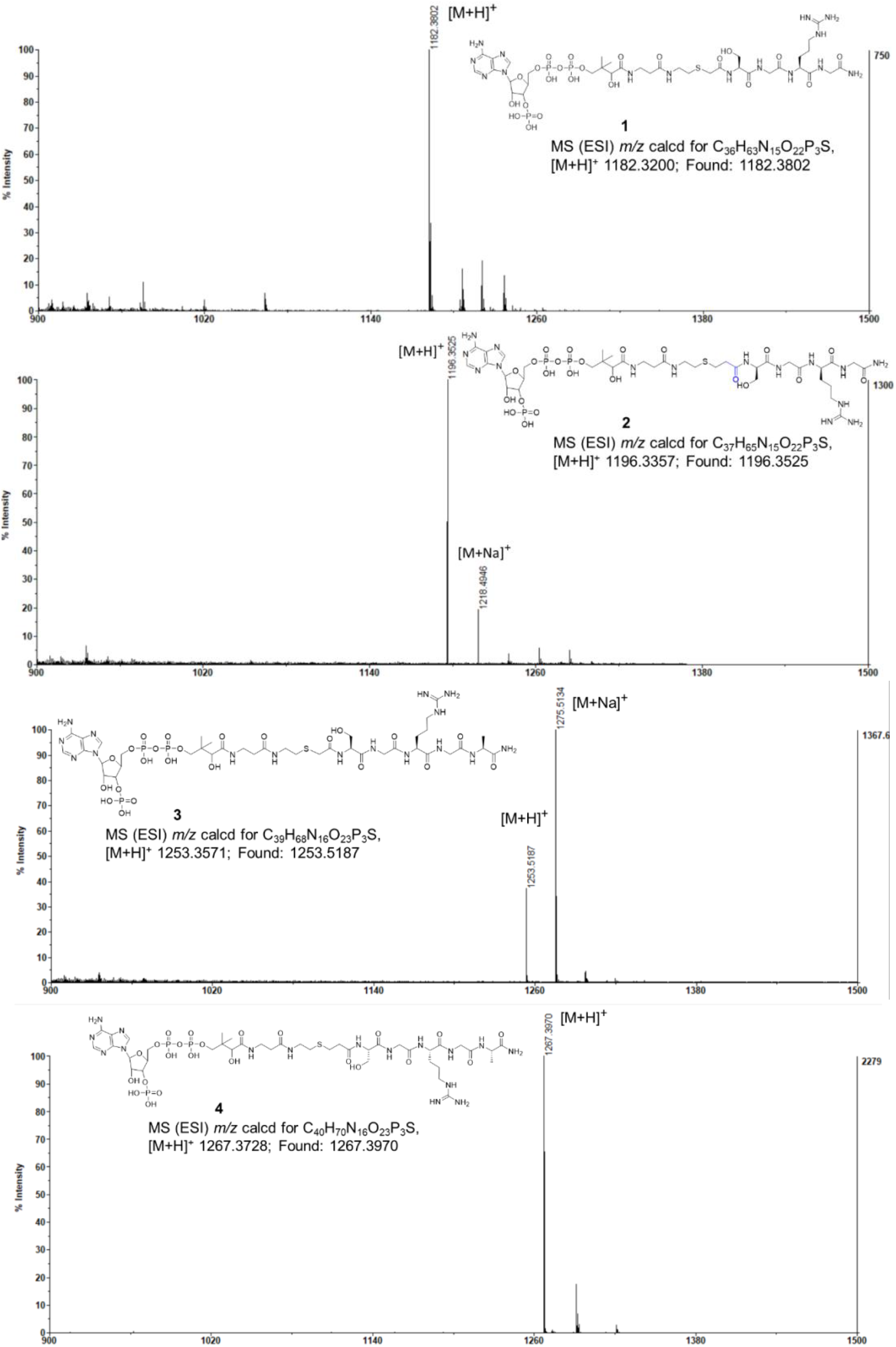
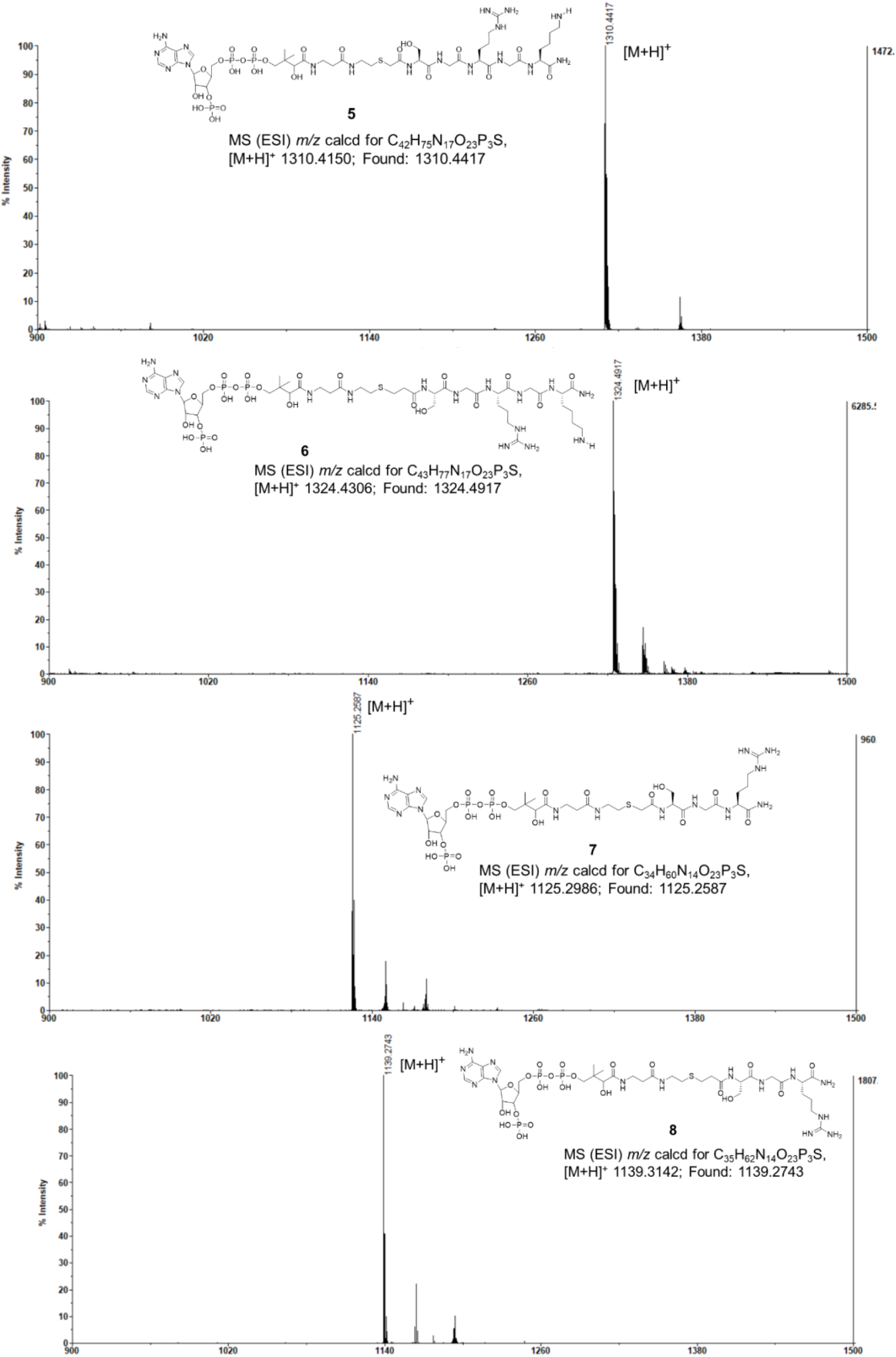
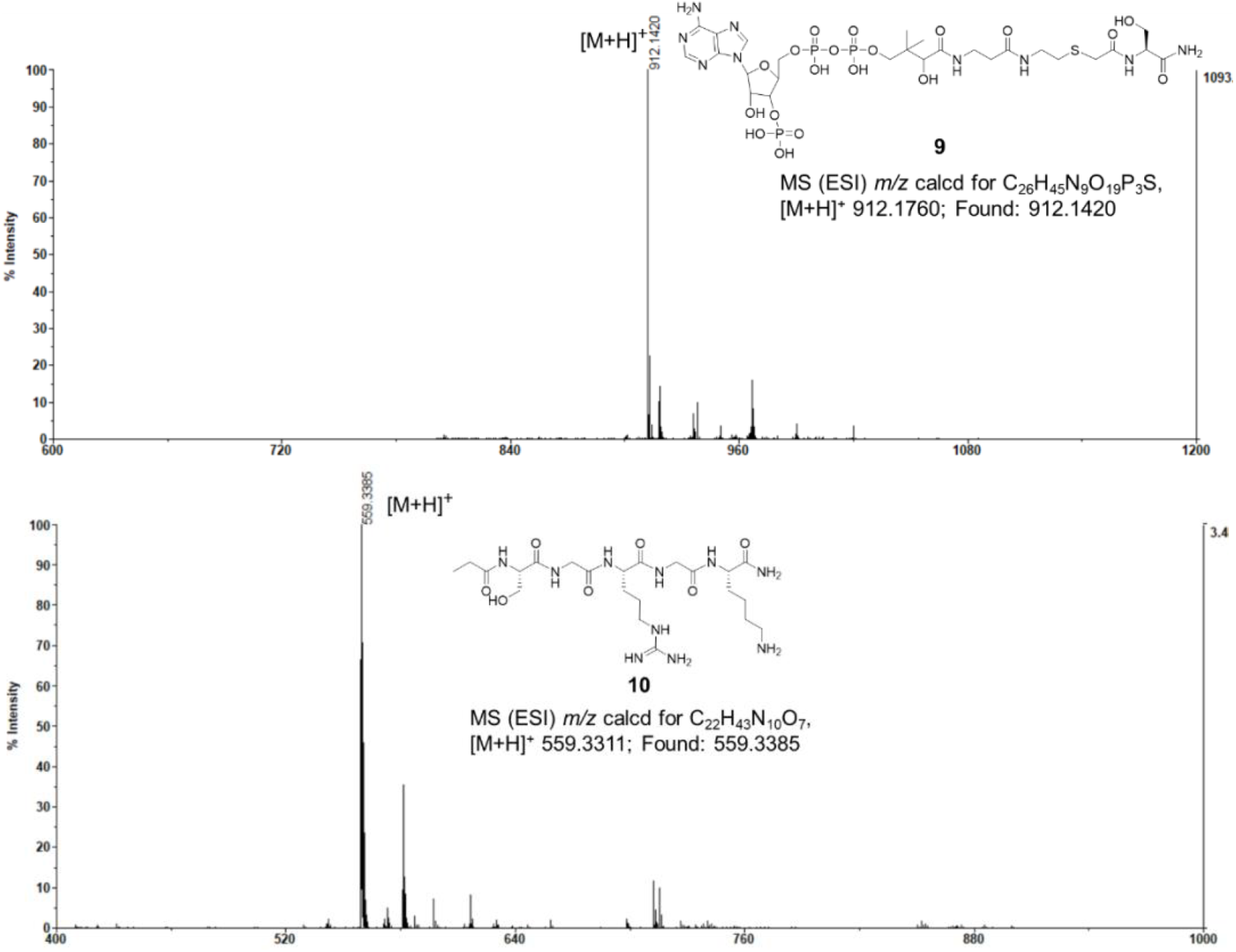
HRMS of Compound 1–9.

**Figure S2.**
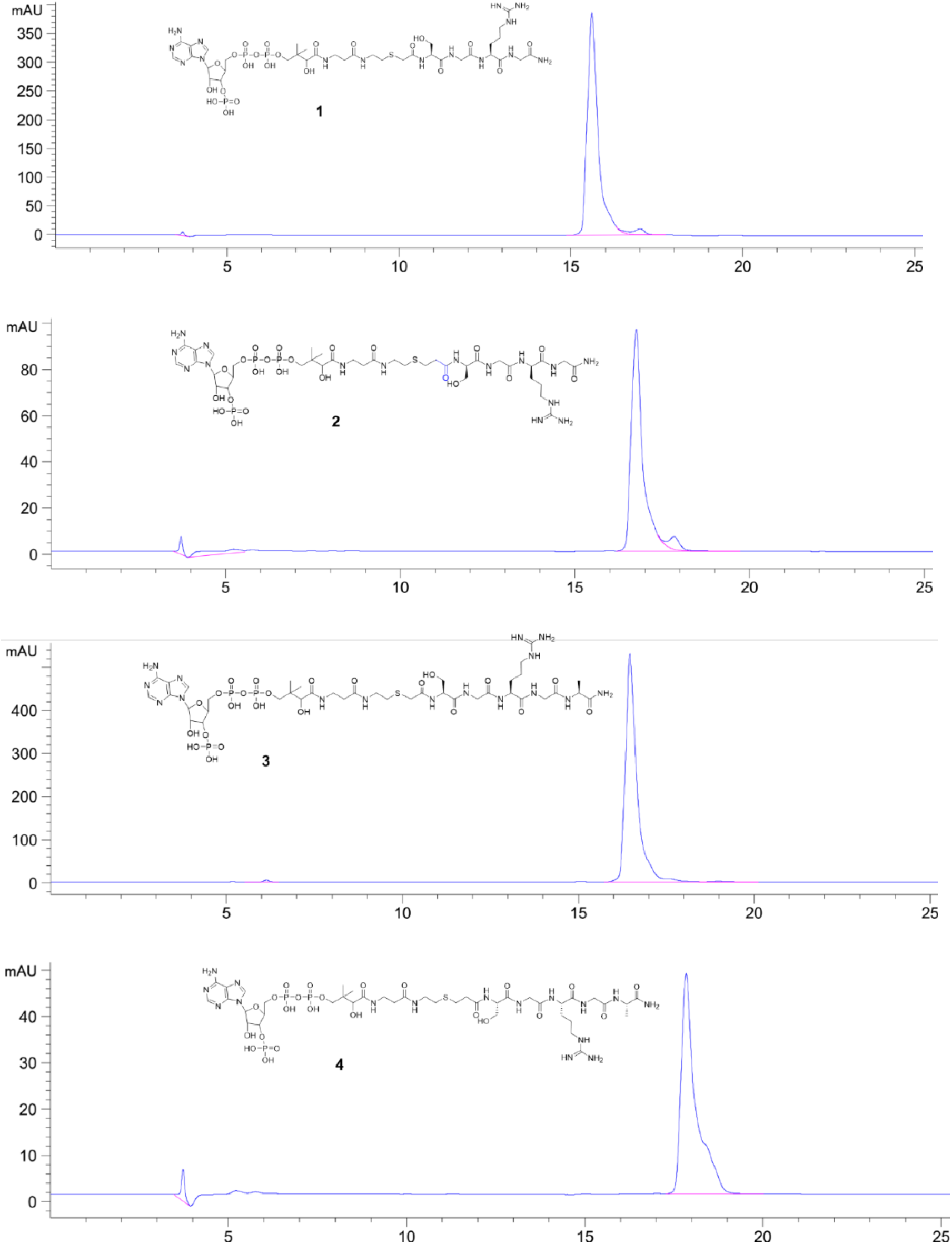
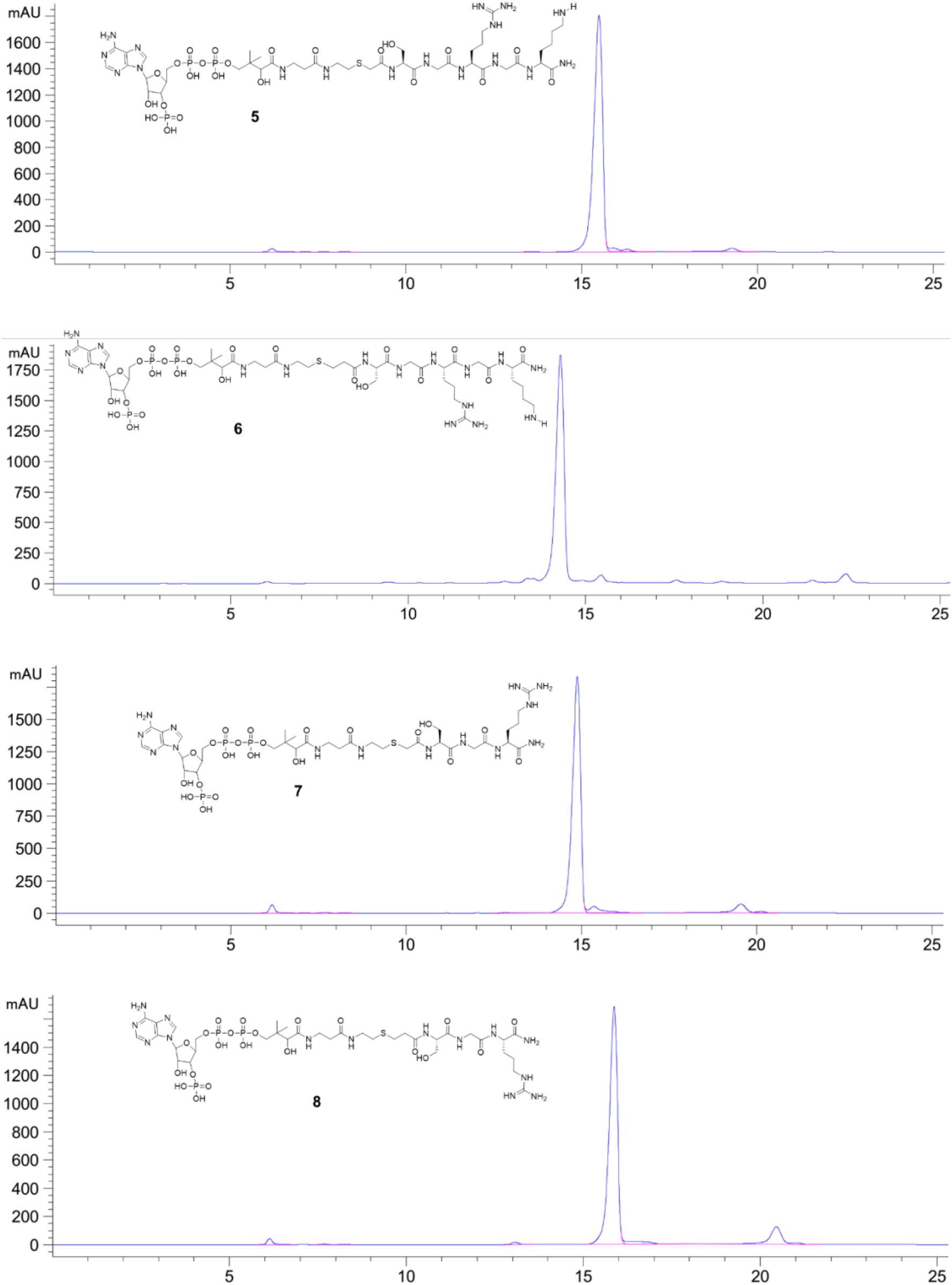
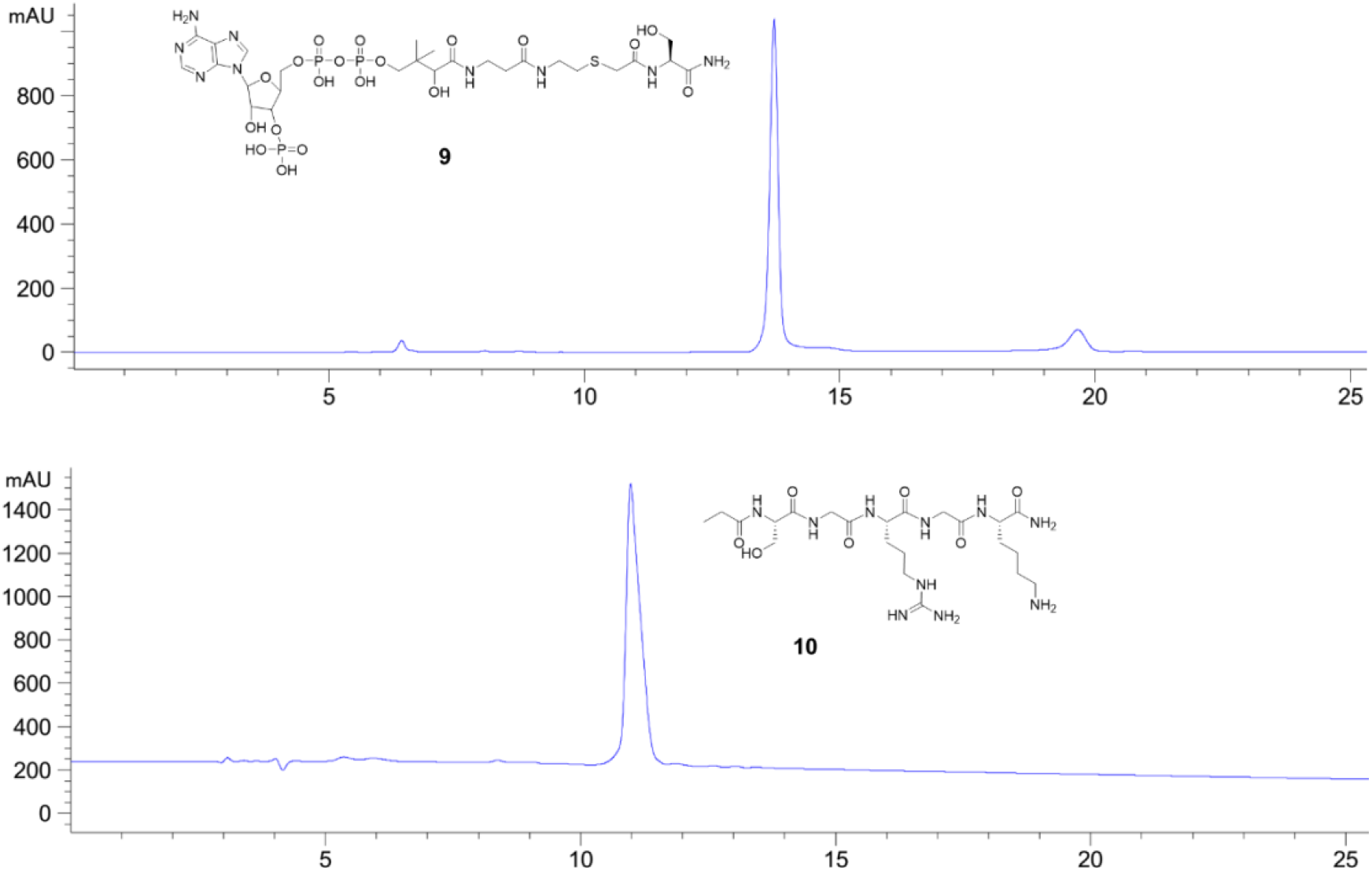
LC Spectrum of Compound 1–9.

**Figure S3.**
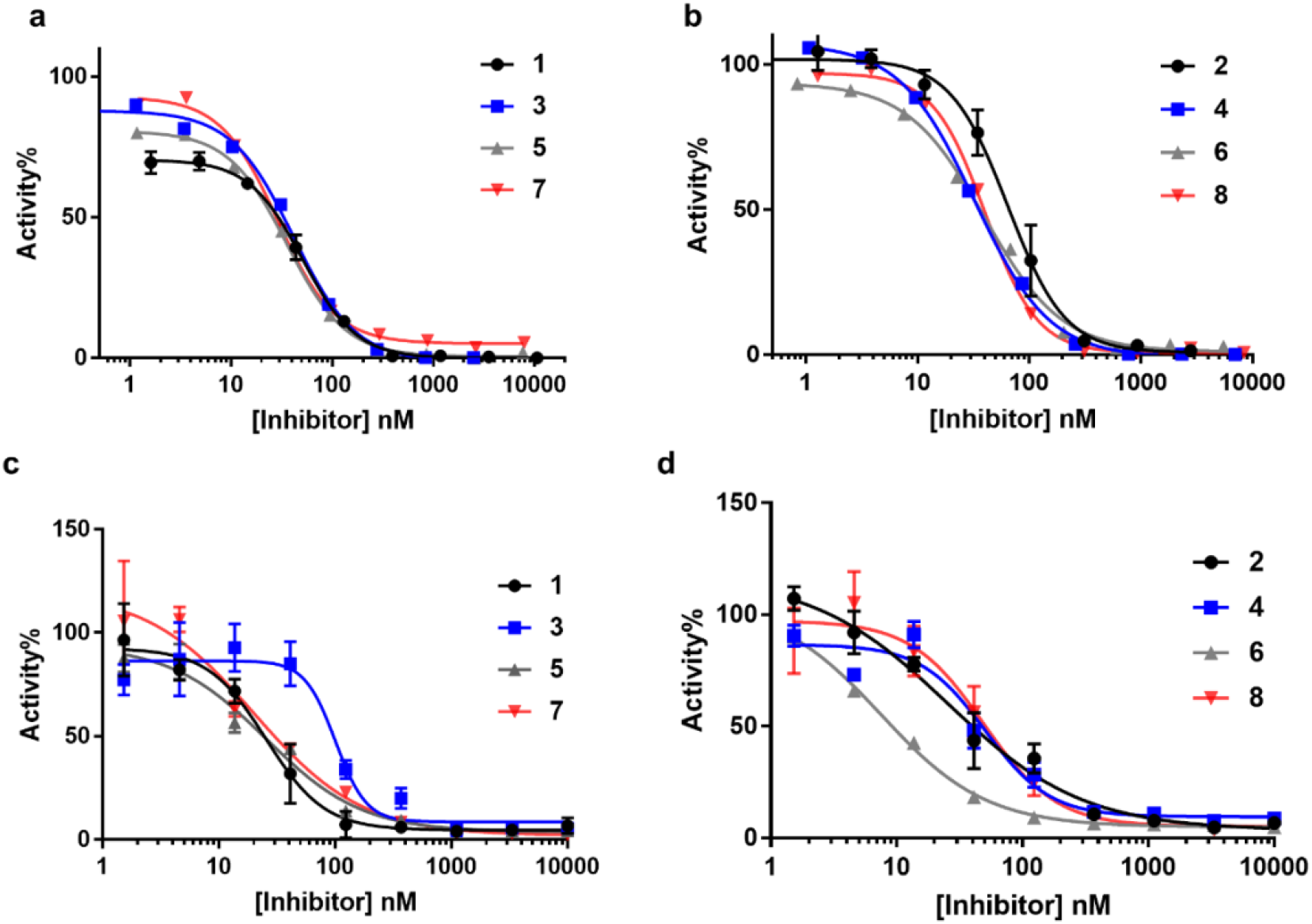
IC_50_ fitting curves of compounds in the fluorescence and radio isotopic assay. a) Concentration–response plots of bisubstrate analogues with an acetyl linker under 1xKm of H4-8 peptide and AcCoA condition (n=2) in the fluorescence assay. b) Concentration–response plots of bisubstrate analogues with a propionyl linker under 1xKm of H4-8 peptide and AcCoA condition (n=2) in the fluorescence assay. c) Concentration-response plots of bisubstrate analogues with an acetyl linker under 4x*K*_m_ of H4-19 peptide and AcCoA condition (n=3) in the radio isotopic assay. d) Concentration-response plots of bisubstrate analogues with a propionyl linker under 4x*K*_m_ of H4-19 peptide and AcCoA condition (n=3) in the radio isotopic assay.

**Figure S4.**
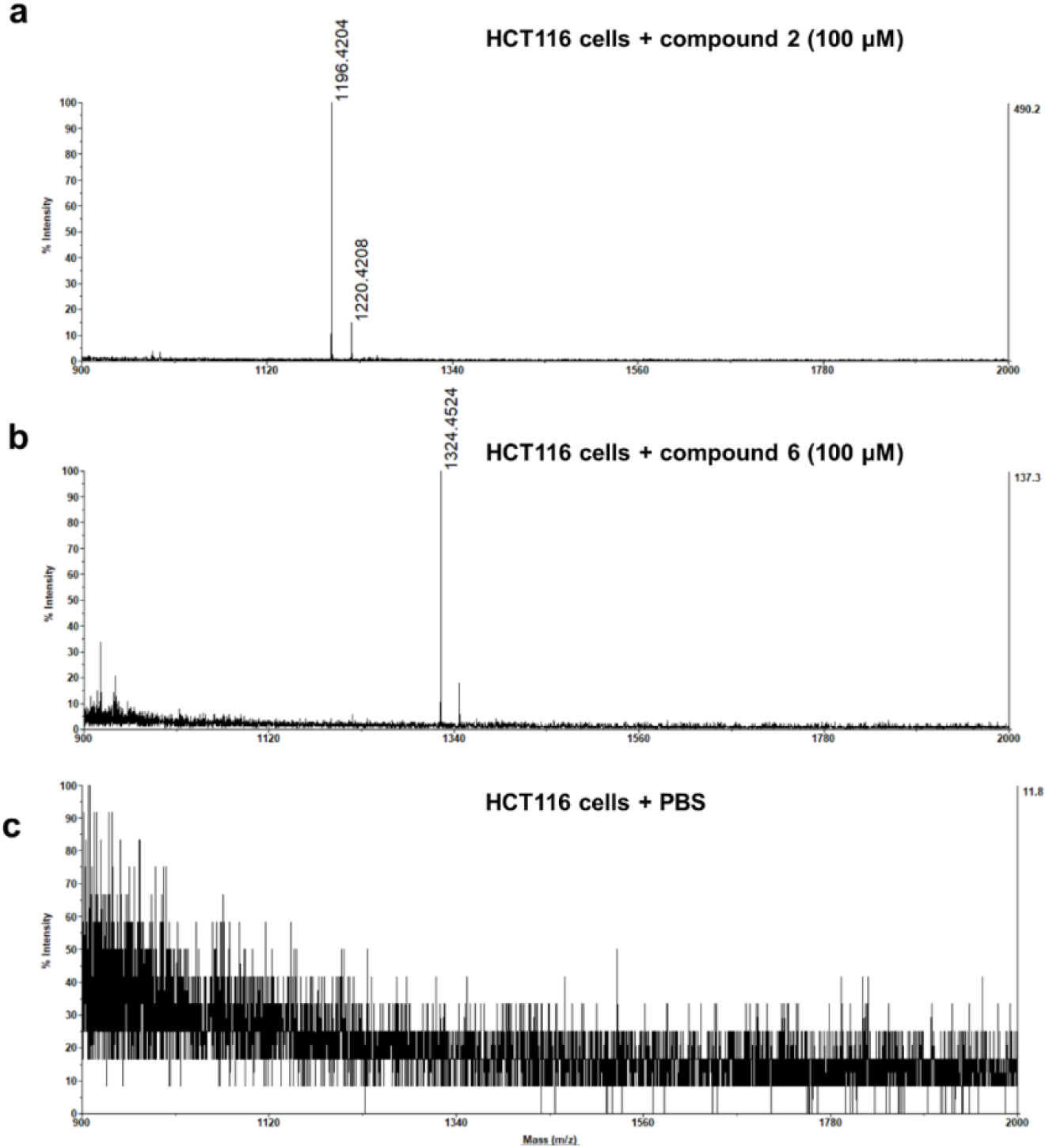
Cell permeability study. a) MALDI-MS for incubation of 100 μM compound **2** with HCT116 cells; **b**) MALDI-MS for incubation of 100 μM compound **6** with HCT116 cells; **c**) MALDI-MS for PBS with HCT116 cells. [**2**+H]^+^ 1196.3357, [**2**+Na]^+^ 1220.4208; [**6**+H]^+^ 1324.4524.

**Table S1.**
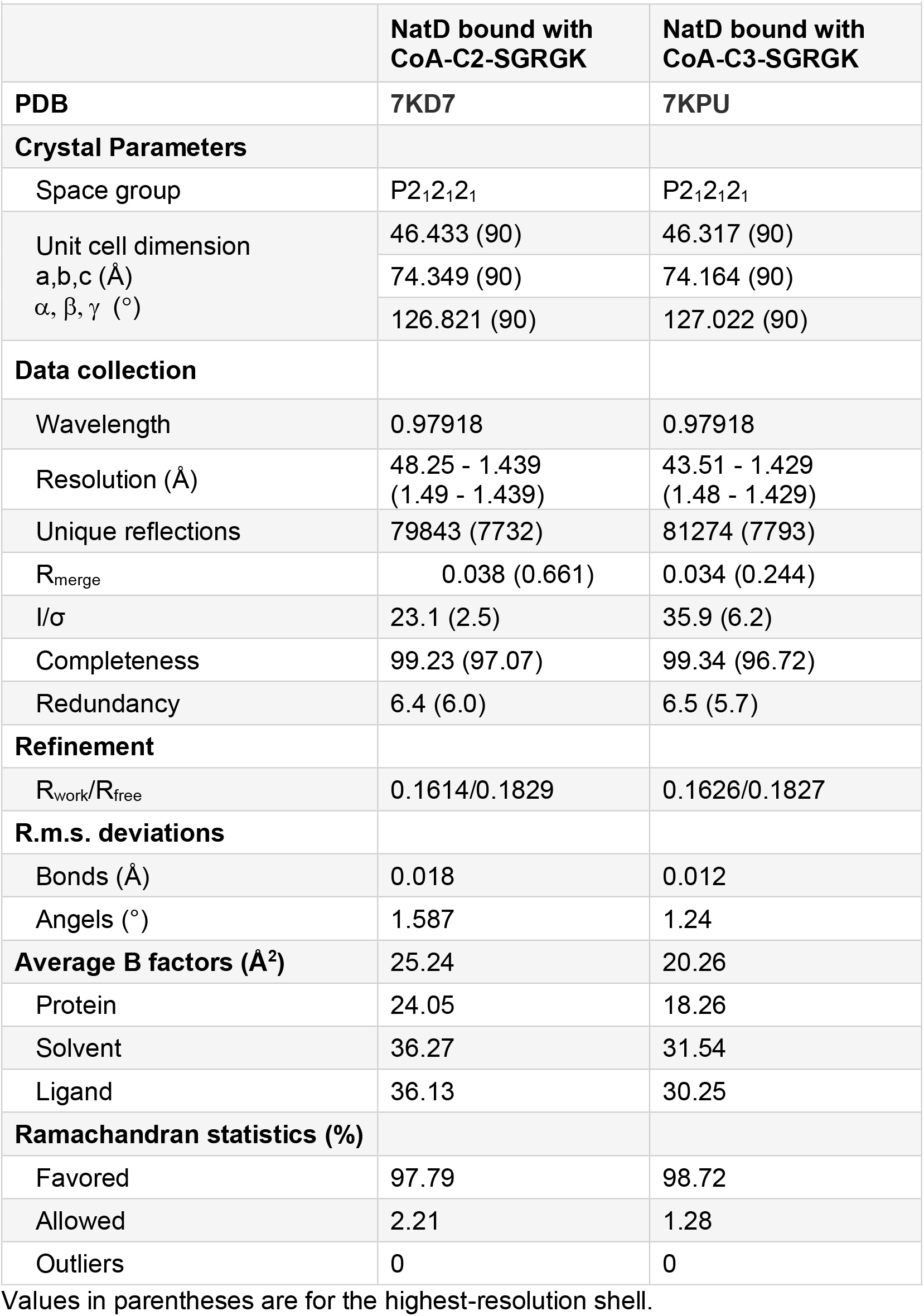
Data statistics for NatD crystal structures in complex with inhibitors.

|  | NatD bound with CoA-C2-SGRGK | NatD bound with CoA-C3-SGRGK |
| --- | --- | --- |
| **PDB** | **7KD7** | **7KPU** |
| **Crystal Parameters** |  |  |
| Space group | P2<sub>1</sub>2<sub>1</sub>2<sub>1</sub> | P2<sub>1</sub>2<sub>1</sub>2<sub>1</sub> |
| Unit cell dimension<br>a,b,c (Å)<br>$\alpha, \beta, \gamma$ (°) | 46.433 (90) | 46.317 (90) |
|  | 74.349 (90) | 74.164 (90) |
|  | 126.821 (90) | 127.022 (90) |
| **Data collection** |  |  |
| Wavelength | 0.97918 | 0.97918 |
| Resolution (Å) | 48.25 - 1.439<br>(1.49 - 1.439) | 43.51 - 1.429<br>(1.48 - 1.429) |
| Unique reflections | 79843 (7732) | 81274 (7793) |
| R<sub>merge</sub> | 0.038 (0.661) | 0.034 (0.244) |
| I/$\sigma$ | 23.1 (2.5) | 35.9 (6.2) |
| Completeness | 99.23 (97.07) | 99.34 (96.72) |
| Redundancy | 6.4 (6.0) | 6.5 (5.7) |
| **Refinement** |  |  |
| R<sub>work</sub>/R<sub>free</sub> | 0.1614/0.1829 | 0.1626/0.1827 |
| **R.m.s. deviations** |  |  |
| Bonds (Å) | 0.018 | 0.012 |
| Angles (°) | 1.587 | 1.24 |
| **Average B factors (Å<sup>2</sup>)** | 25.24 | 20.26 |
| Protein | 24.05 | 18.26 |
| Solvent | 36.27 | 31.54 |
| Ligand | 36.13 | 30.25 |
| **Ramachandran statistics (%)** |  |  |
| Favored | 97.79 | 98.72 |
| Allowed | 2.21 | 1.28 |
| Outliers | 0 | 0 |

Values in parentheses are for the highest-resolution shell.

## Notes

### Competing Interest Statement

The authors have declared no competing interest.

### Summary of Updates

Add inhibition mechanism and cellular target engagement.

